# Microsecond resolved infrared spectroscopy on non-repetitive protein reactions by applying caged-compounds and quantum cascade laser frequency combs

**DOI:** 10.1101/2021.01.04.425172

**Authors:** Mohamad Javad Norahan, Raphael Horvath, Nathalie Woitzik, Pierre Jouy, Florian Eigenmann, Klaus Gerwert, Carsten Kötting

## Abstract

Infrared spectroscopy is ideally suited for the investigation of protein reactions at the atomic level. Many systems were investigated successfully by applying Fourier transform infrared (FTIR) spectroscopy. While rapid-scan FTIR spectroscopy is limited by time resolution (about10 ms with 16 cm^-1^ resolution), step-scan FTIR spectroscopy reaches a time-resolution of about 10 ns but is limited to cyclic reactions that can be repeated hundreds of times under identical conditions. Consequently, FTIR with high time resolution was only possible with photoactivable proteins that undergo a photocycle. The huge number of non-repetitive reactions, e.g. induced by caged compounds, were limited to the ms time domain. The advent of dual comb quantum cascade laser allows now for a rapid reaction monitoring in the μs time domain. Here we investigate the potential to apply such an instrument to the huge class of G-proteins. We compare caged-compound induced reactions monitored by FTIR and dual comb spectroscopy, respectively, by applying the new technique to the α subunit of the inhibiting G_i_ protein and to the larger protein-protein complex of Gα_i_ with its cognate regulator of G-protein signaling (RGS). We observe good data quality with 4 μs time resolution with a wavelength resolution comparable to FTIR. This is more than three orders of magnitude faster than any FTIR measurement on G-proteins in the literature. This study paves the way for infrared spectroscopic studies in the so far unresolvable μs time regime for non-repetitive biological systems including all GTPases and ATPases.

## INTRODUCTION

FTIR spectrometers revolutionized infrared spectroscopy in the 70s of the last century.^1^ Likely, the advent of stable mid-IR quantum cascade lasers (QCLs) will impact infrared spectroscopy to the same extent. Conventional QCLs are superior with regard to brilliance,^2^ but they lack the multiplex advantage of FTIR that is especially helpful in time-resolved measurements of proteins. For systems that can be excited repetitively, step-scan FTIR can provide time-resolved spectra with ns resolution.^3^ For these repetitive systems, tunable QCLs provide time resolved spectra with 10 ns.^2,4^ Pump-probe experiments (vis-pump and IR probe) even allow for femtosecond time-resolved IR spectroscopy.^5^ However, samples that allow only single excitations can either be measured only at a single wavelength^6^ or with rapid-scan FTIR.^7^ The time-resolution of rapid-scan FTIR depends on the scanning velocity of the Michelson interferometer and is limited to about 10 ms at a wavelength resolution of 16 cm^−1^ within high end research FTIR instruments. The implementation of a faster Michelson interferometer allows, in principle, for a higher time resolution, but with a conventional globar as the light source, signal to noise ratios (S/N) are not sufficient for single shot experiments on biological systems.^8^ Synchrotron irradiation and a dispersive setup in combination with an array detector permits single shot experiments with µs resolution.^9^However, even here S/N was not sufficient for a single shot experiments of a protein reaction.^10^

The recent development of dual comb QCL based spectrometers allows for µs time resolution as well, but with a much smaller footprint.^11,12^ In these instruments two broad band lasers that emit at many discrete wavelengths are used as the light source (Figure 1). The wavelength spacing of the first laser (*f*_*rep,1*_) is close but not identical to the other laser (*f*_*rep,2*_). Overlaying the two lasers produces a set of beatings spaced by Δ *f*_*rep*_=*f*_*rep,1*_-*f*_*rep,2*_ in the radiofrequency domain measured by a high bandwidth MCT detector. From this measurement the entire heterodyne beating pattern can be recovered. In our setup Δ *f*_*rep*_ leads to a time resolution of 4 µs. The spectral window of the lasers is from 1207.0 cm^-1^ to 1276.8 cm^-1^. By guiding the QCLs through a sample that is irradiated by a pulsed UV-laser, UV-light induced spectral changes of the sample can be obtained with the time resolution of 4 µs.^11^

**Figure 1.**
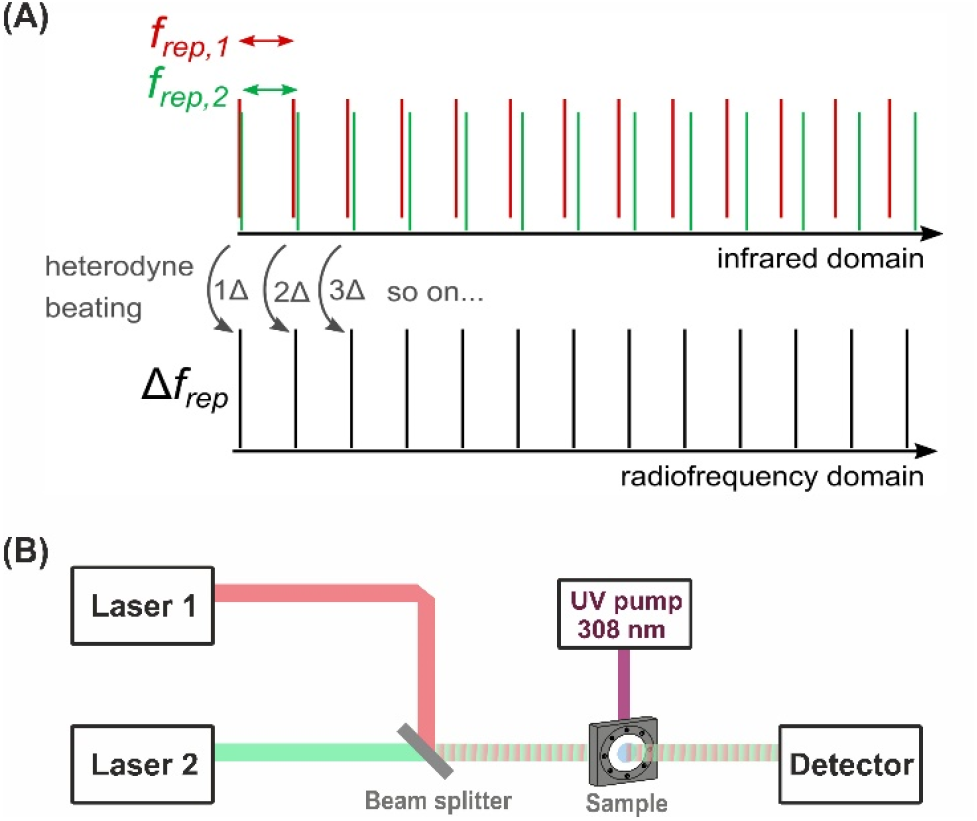
(A) Principle of dual comb spectroscopy. (B) Schematic of the setup of the dual-comb spectrometer used here.

Caged compounds^13^ are a tool for the investigation of reactions of biological systems that can otherwise not be initiated by light. Caged compounds release the reactive compound upon irradiation and cannot be excited repetitively. Rapid-scan FTIR spectrometers usually have a time resolution of about 10 ms at 16 cm^-1^ spectral resolution. Due to limited S/N the actual time-resolution for measuring protein reactions is often even slower and often artificial low-temperatures are necessary to slow down the reaction. Most prominently ATPases^14^ and GTPases^15^ were investigated in detail by means of caged nucleotides. As caged compounds the P^3^-[1-(2-nitrophenyl)ethyl] ester (NPE)^16^ and the P^3^-[para-hydroxyphenacyl] ester (pHP)^17^ of the nucleotides were used.

Here we used, for the first time, a dual comb QCL based spectrometer for the investigation of GTPases in the µs time-domain. First, we measured the photolysis reaction the two caged-GTPs, NPE-GTP^16^ and pHP-GTP^18^, in solution without a protein present and compare the results with conventional rapid-scan FTIR. In the next step we applied the technique to the Gα subunit of the heterotrimeric G_i_ protein and monitor its GTPase reaction.^19^ Finally, the RGS catalyzed reaction of Gα_i_ was measured for the first time at ambient temperatures by time-resolved infrared spectroscopy. At room temperature, the reaction is completed before the first datapoint of a rapid-scan FTIR measurement can be recorded. With dual comb IR it is well resolved, including a so far unknown intermediate.

## EXPERIMENTAL SECTION

The P^3^-[1-(2-nitrophenyl)ethyl] ester (NPE) of GTP was obtained from Jena Bioscience (Jena, Germany). P^3^-[para-hydroxyphenacyl] ester (pHP) of GTP was synthesized by coupling GDP with pHP-caged P_i_. pHP-P_i_ was obtained in five steps from para-hydroxy-acetophenone and dibenzylphosphate.^17^

Gα_i1_ and RGS proteins were expressed and purified as described by Mann et al..^19^ In the purified proteins the nucleotide GDP was exchanged for pHP-GTP. The exchange rate was >95% as checked by reversed phase HPLC (LC-2010; Shimadzu) [mobile phase: 50 mM Pi (pH 6.5), 5 mM tetrabutylammoniumbromide, 7.5% (vol/vol) acetonitrile; stationary phase: ODS-Hypersil C18 column]. For the intrinsic Gα_i1_ measurements Gα_i1_ pHP-GTP was lyophilized. For the samples, lyophilized protein was resuspended in buffer to reach the following concentrations: 5mM Gα_i1_, 200 mM Hepes (pH 7.5), 150 mM NaCl, 5 mM MgCl_2_, 200 mM DTT, and 0.1% (vol/vol) ethylene glycol. For the RGS catalyzed reactions a 1:1 molar ratio of Gα_i1_ with RGS was lyophilized and resuspended to reach 5mM Gα_i1_·RGS, 100 mM Hepes (pH 7.5), 100 mM Tris (pH 7.5), 150 mM NaCl, 5 mM MgCl_2_, 20 mM DTT, and 0.1%(vol/vol) ethylene glycol. The samples were packed between two CaF_2_ windows, separated by a spacer ring yielding a pathlength of about 60 µm, that were sealed with silicon grease and mounted either in a Bruker Vertex80v spectrometer or an IRsweep IRis-F1 dual-comb spectrometer. All measurements were done at room temperature (293 K). The reactions were initiated by flashes of a XeCl-excimer laser (308 nm, 150 mJ, Coherent LPX Pro 240). FTIR measurements were recorded at 4 cm^-1^ spectral resolution, manipulated by zero filling by a factor of 2, and Fourier-transformed using Mertz phase correction and Blackman–Harris three-term apodization function.

The operating principles of the IRis-F1 spectrometer have been described in detail.^11,12,20^ Briefly, two continuous wave QCL frequency combs with Δ*f*_*rep*_ = 3.2 MHz were used to obtain the heterodyne beating signal. At the operating conditions used here, *f*_*rep,1*_ and *f*_*rep,2*_ were about 9.82 GHz, as determined from the inter-line beating measured with a spectrum analyzer (FSW26, Rhode & Schwarz). This gives a native spectral resolution of 0.3275 cm^-1^. The spectral output of the individual lasers is presented in Figure S1. The power at the sample was adjusted to be about 3 mW. We obtained data in two measurement modes, as described further in Figure S2. In time resolved mode, each acquisition is 33.6 ms long, sampled at 2 GS/sec, and fast Fourier transforms are taken at intervals of 4 µs, which determines the time resolution of the measurement. Time resolutions up to 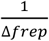 are possible,^11,21^ but given the rates of the processes investigated here, and a drop in signal to noise at lower integration times, 4 µs was chosen. In this mode, each additional acquisition takes a relatively long time to complete and serves only to improve the signal to noise ratio by synchronizing with repeated excitations. In long-term mode, each acquisition is 1 ms long and repeats with a frequency of approximately 100 Hz. The time-resolved dual-comb IR data were obtained from combining measurements from both modes, which were each logarithmically averaged. Before merging, the data were scaled according to the difference spectra at peaks wavenumber of each dataset. Furthermore, to avoid discontinuities, we allowed for ∼15 ms overlap between both data sets. The originally recorded high spectral resolution of the IRsweep instrument at 0. 3275 cm^-1^ was also averaged to 4.2 cm^-1^ for better S/N.

The data was further analyzed by a global fit (Eq.1).^15^ The time-resolved absorbance change Δ*A*(*ν,t*) is described by the absorbance change induced by photolysis *a*_ph_(*ν*) followed by a number n of exponential functions fitting the amplitudes *a* for each wavenumber ν. The amplitude spectra have negative peaks for absorptions that disappear (or lose intensity) and positive peaks for absorptions that emerge (or gain intensity).

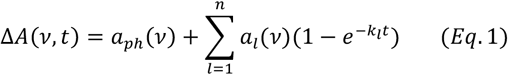

For easier comparison, we give the half-life (t_1/2_=ln(2)/k) in the manuscript. Due to the limited intensity of the excitation beam and the insufficient overlap of the excitation laser with the absorption spectrum of the caged-compounds, not the complete sample is photolyzed by one laser flash. Note that a more intense excitation beam could also lead to an increased heat artifact. To estimate the amount of photolysis the signal strength at the most intense analyte peak (1253 cm^-1^) was integrated over the first 2 ms after excitation as a function of sample excitation number. This was repeated for the negative times (−2 to 0 ms) as a control, as no spectral features are expected there. Figure S3 indicates that for NPE-GTP after 20 excitations, no further photolysis occurs. For pHP-GTP after 10-15 excitations, no further photolysis is observed (Figure S4) and the data shown are the averages of the first 10 excitations. In the Figures of the main text, we always show the results of measurements obtained by a single sample.

## RESULTS AND DISCUSSION

Photolysis of NPE-GTP does not produce GTP directly, first an intermediate, the aci-nitro anion is formed (Figure 2A).^22^ Depending on the reaction conditions, GTP is formed from this intermediate, usually in the ms time regime. We measured NPE-GTP photolysis with both a Bruker Vertex 80v FTIR spectrometer and the IRsweep IRis-F1 dual-comb spectrometer (Figure 2).

**Figure 2.**
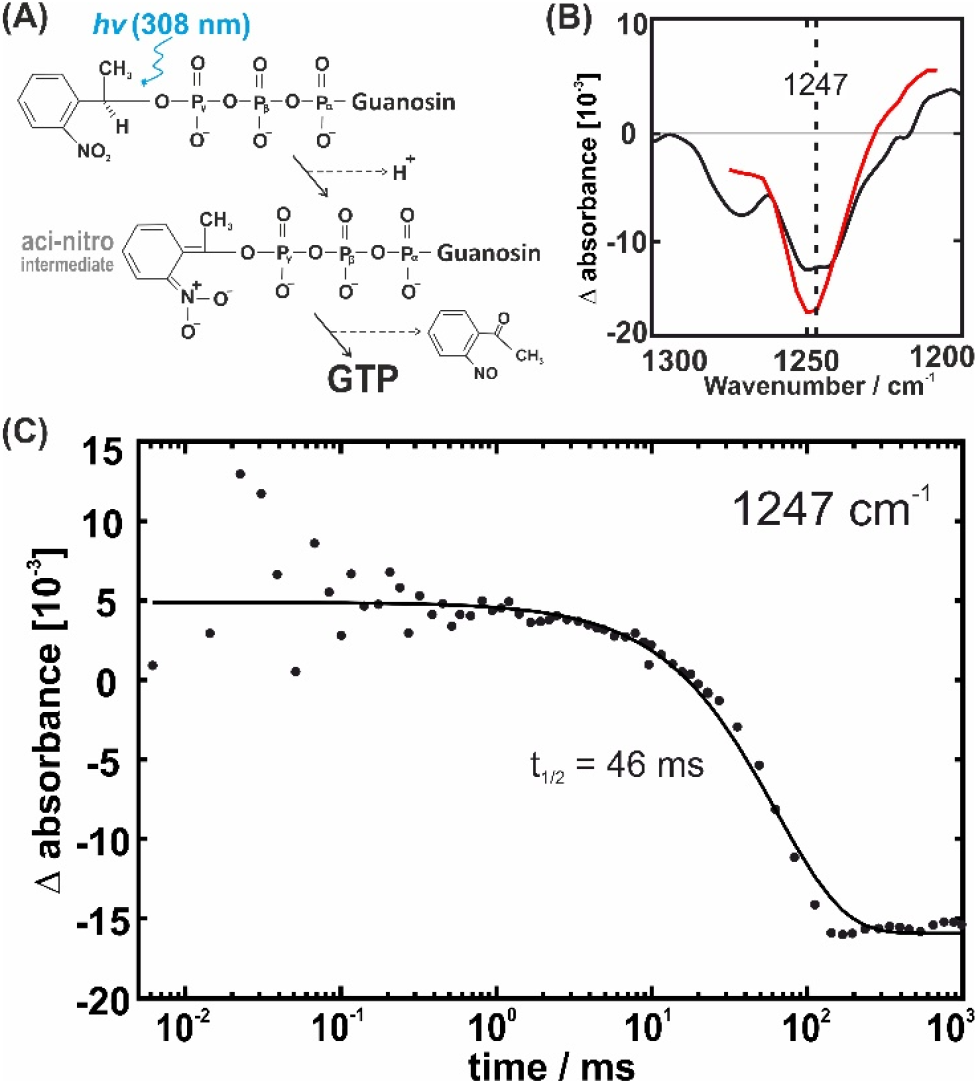
(A) Photolysis of NPE-GTP Reaction Scheme. (B) Photolysis spectra obtained by FTIR (black) and dual-comb (red) obtained by global fits of the time-resolved data, respectively. (C) Kinetics of the hydrolysis reaction obtained by dual-comb experiments. The points correspond to the time-resolved data and the line to the global fit according to equation 1.

The photolysis difference spectrum of the FTIR experiment can be compared with the same difference from the dual-comb experiment. In both cases the spectra are shown in a way that the newly formed absorptions are facing upwards and vanishing absorptions are facing downwards. Figure 2B shows that the spectra agree nicely. The spectral window of the dual comb experiment shows nicely the vanishing band of the combined asymmetric stretching vibrations of the phosphate groups of the aci-nitro-GTP intermediate.^23^

The same reaction was investigated before by a step-scan FTIR-experiment with 10 µs time resolution at a spectral resolution of 15 cm^-1^.^24^ The data were obtained within 5 hours of measurement time, using 200×200 µm^2^ segments from five samples of 1 cm^2^ area.^25^ In comparison, the dual comb experiment was done with a single sample. Compared to the 5 hours of the step-scan FTIR experiment the dual comb measurement only took a few seconds. Further, much smaller amounts of sample are needed. The saving in sample consumption is about a factor of 10 in our experiment but could be improved by another factor of 10 by using a cuvette optimized for the QCL profile. The diameter of the laser beam is about 3 mm and much smaller than the conventional IR cuvettes as shown in Figure S5. The quantum yield of NPE-GTP hydrolysis with our 308 nm XeCl-excimer laser is limited. For this reason, we repeated the experiment 20 times with the same sample in the same position and coadded the observed changes. The sample response after each shot was monitored (Figure S3). The kinetics in Figure 2B were obtained by combining the time resolved and long term measurement modes of the IRis-F1 (details of the data treatment are given in the Methods section). We measured a half-life of 46 ms at 293 K for the intermediate.

pHP-GTP is the superior caged compound for the investigation of fast reactions because photolysis is fast and without an intermediate. Indeed, we observe very fast production of GTP from pHP-GTP with only one minor and very fast kinetic rate that might be a heat artefact (Figure S4). For this reason, we use pHP-GTP for the protein reactions shown below.

The protein with its GTPase domain (orange) and all-alpha domain (yellow) is shown in Figure 3A. The central nucleotide is shown in a ball and sticks representation. Figure 3B shows the reaction scheme of Gα_i_. After irradiation with the 308 nm excimer laser, we expect the photolysis reaction and subsequently the hydrolysis. The hydrolysis reaction is slow and can be observed by rapid-scan FTIR as a control for our first dual comb experiments with a protein. In both cases a global fit of the time-resolved data by applying equation 1 with n=1 describes the data well. This can be recognized for the DCS experiment by the nice agreement of the time resolved data at 1215 cm^-1^ with the curve from the fit. Comparing DSC and FTIR, the amplitude spectra of photolysis a_ph_ and hydrolysis spectra a_1_ both agree nicely (Figure 3C&3D). The absorption peaks were assigned in the literature by isotopic labelling.^19,26^ In the photolysis a combined asymmetric stretching vibrations of the phosphate groups is assigned and in the hydrolysis the asymmetric PO_2_ stretching mode of the α-phosphate of GTP is shown. Further, the single exponential kinetics (Figure 3E) is in line with the literature.^19^ After having demonstrated the basic functioning of the dual comb technique for GTPase reactions of proteins, we want to investigate a very fast reaction, which cannot be observed at room temperature by FTIR. The RGS catalyzed reaction of Gα_i_ is much faster than the reaction of Gα_i_ alone and cannot be resolved by rapid-scan FTIR at ambient temperature. The first data-point in the FTIR measurements of these larger protein-protein complexes (Figure 4A) is usually above 100 ms (see e.g. Figure 2B in ^19^). With the dual comb experiment our first datapoint is at 4 µs. The kinetics at 1240 cm^-1^ show nicely the decay of the α-GTP band due to the GTPase reaction.^19^ Clearly, the reaction is almost completed at 100 ms, indicating that the reaction could not be observed by rapid scan FTIR at all. A half-life of 90 ms was obtained. Interestingly there are even two additional very fast rates (Figure 4C half-lives of 38 µs and 86 µs) resolved, preceding hydrolysis. We can speculate that one rate corresponds to a heating artifact (as observed for pHP-GTP alone, Figure S3) and the other to a fast rearrangement within the catalytic site. Such a rearrangement was also observed on a slower timescale in a Gα_i_ mutant.^27^ However, further experiments including the measurements of further Gα_i_ mutants will be necessary for a clear-cut assignment and is not within the scope of this work. The complete reaction with all the information obtained in the dual comb experiment is shown in Figure 4D.

**Figure 3.**
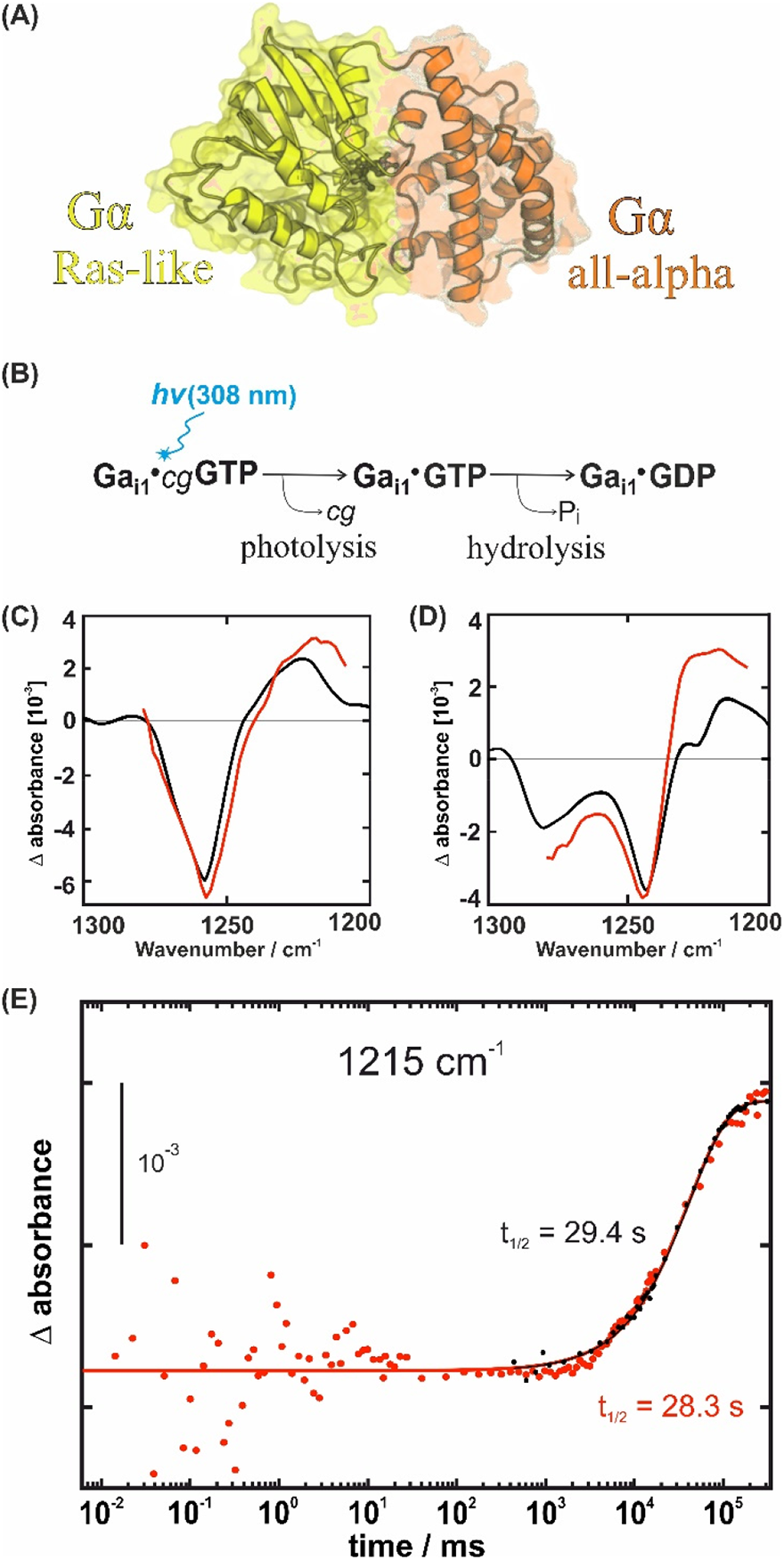
(A) Structural model of Gai from pdb ID 2G83. (B) Reaction scheme for the GTPase reaction of Gα_i_. (C) Photolysis and (D) hydrolysis spectra obtained by FTIR (black) and dual-comb (red) obtained by global fits of the time-resolved data, respectively. (E) Kinetics of the hydrolysis reaction. The points correspond to the time-resolved data and the lines to the global fits according to equation 1. The absorption of the dual comb experiment was normalized to the FTIR experiment, where a thinner sample was used.

**Figure 4.**
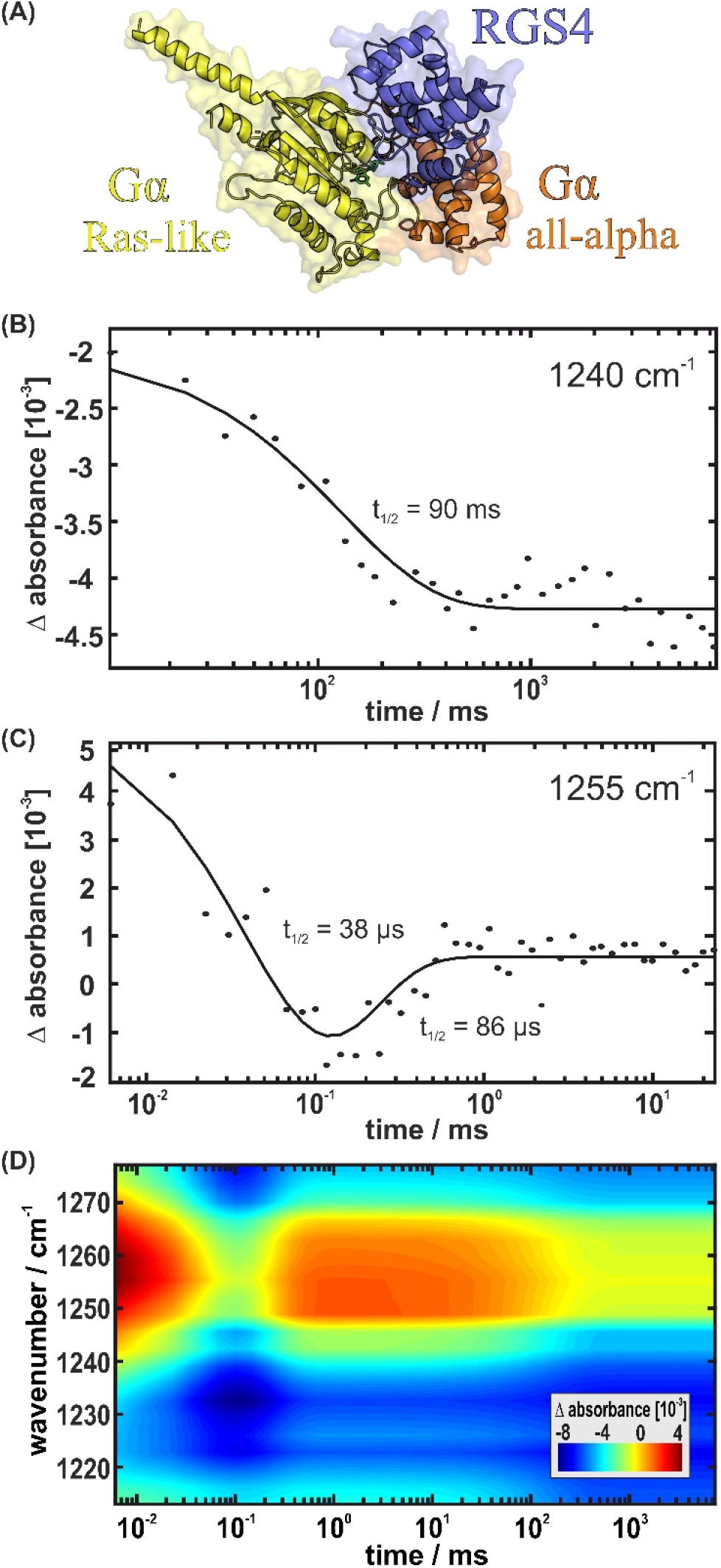
(A) Structural model of Gαi·RGS from pdb ID 1AGR. (B) Kinetics of the hydrolysis reaction of Gαi·RGS. (C) Additional pre-hydrolysis rates obtained by dual comb IR. In (B) and (C) the points correspond to the time-resolved data and the line to the global fit according to equation 1. (D) 3D-plot of the changes obtained from the global fit (equation 1) of the dual comb data as detailed in the methods section. After photolysis three processes (n=3) with half-lives of 38 µs, 86 µs and 90 ms were obtained.

## CONCLUSIONS

Overall, we were able to show that the dual comb setup is very suitable for the investigation of proteins with caged compounds. The only drawback of the new technique is the relatively small spectral window of each dual comb setup. However, laser modules can be changed and modules for all interesting wavelengths between 2200-900 cm^-1^ are available and with the about 100 times stronger source power single shot analyses of weak absorbers are possible. Another approach could be the measurement of full spectra by FTIR with low time resolution and subsequent measurement of interesting regions with a dual comb setup.

We demonstrate that with a single sample, a time resolution in the µs regime can be obtained even for a larger protein-protein complex. In our setups we use a sample thickness of 60 µm, a great advantage of the intense QCLs in comparison with FTIR, where we use about 10 µm sample thickness. This alone should lead to an about 6 times larger S/N ratio. The larger pathlength also allows for a much easier implementation of flow through setups using conventional microfluidics.

## ACKNOWLEDGMENT

We thank Dr. Daniel Mann for the preparation of Gαi and Dr. Christian Teuber, Adrian Höveler and Kristin Labudda for help with the sample preparation. We further thank Dr. Jonas Schartner for the synthesis of pHP-GTP This work was supported by Deutsche Forschungsgemeinschaft (DFG, German Research Foundation), individual Research Grant GE 599/20-1 to KG and KO 3813/1-1 to CK and Research Training Group 2341 ‘MiCon’. Further support was provided by the Ministry for Culture and Science (MKW) of North Rhine-Westphalia (Germany) through grant 111.08.03.05-133974 and the Protein Research Unit Ruhr within Europe (PURE) funded by the Ministry of Innovation, Science and Research (MIWF) of North-Rhine Westphalia (Germany).

## ASSOCIATED CONTENT

### Supporting Information

The Supporting Information is available free of charge on the ACS Publications website.

Figure S1: Spectral coverage and intensities of the individual QCL devices.

Figure S2: Slow and fast mode of the dual comb instrument. Figure S3: Photolysis signal development during an experiment.

Figure S4: Photolysis of pHP-GTP.

Figure S5: Comparison of the conventional IR cuvette with the beam diameter of the QCL.

