## Supplemental Information for "Microsecond resolved infrared spectroscopy on non-repetitive protein reactions by applying caged-compounds and quantum cascade laser frequency combs"

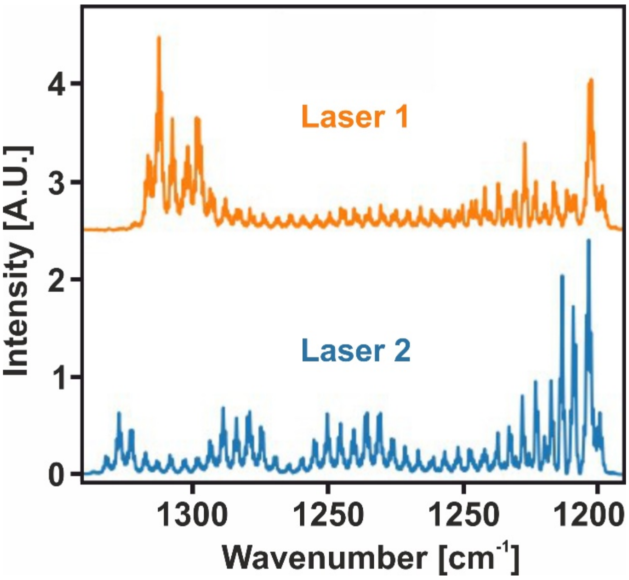


*Figure S1. Spectral coverage and intensities of the individual QCL devices recorded with a Vertex 70 (Bruker) FTIR spectrometer.*

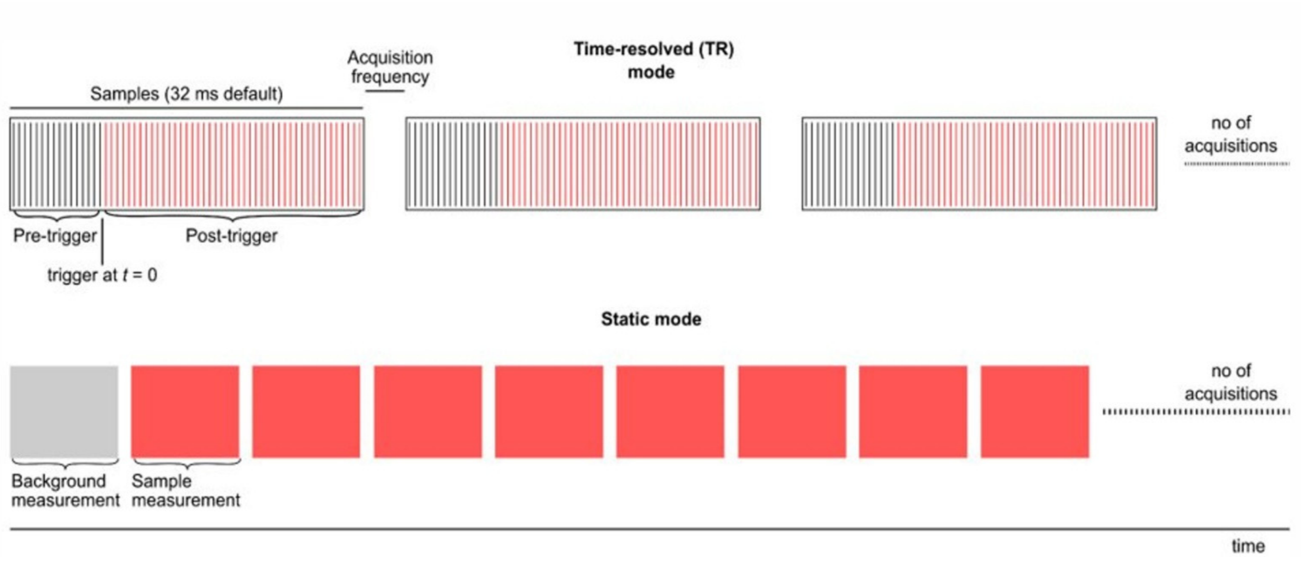


*Figure S2. Illustration of the acquisition modes of the dual comb spectrometer. In the time resolved mode (upper panel) the background and sample data are obtained in the same acquisition, at a rate of 4 μs per spectrum, for a duration of up to ca. 32 ms. The trigger coincides with an external event and delimits background and sample parts. Successive acquisitions are co-added to improve signal to noise. In long the term mode (lower panel), entire acquisitions are integrated into single spectra. Here, the time resolution is given by the acquisition frequency of the measurement.*

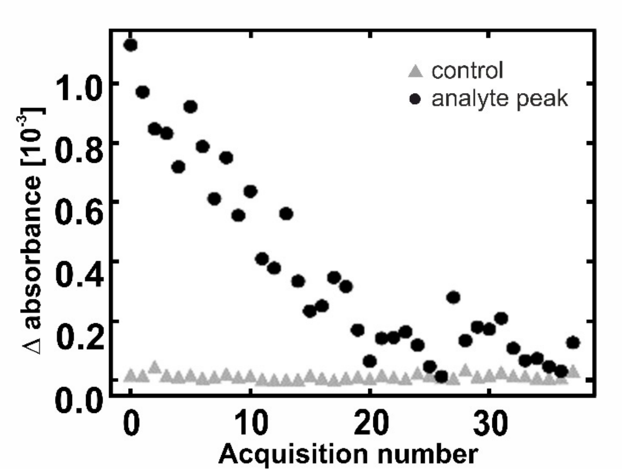


*Figure S3. The magnitude of the spectral response to exciation as a function of excitation pulses. After each flash, the amount of NPE-GTP decreases, leading to a decrease of signal after each pulse. After about 20 pulses all NPE-GTP is consumed. With a better overlap of absorption spectrum of the caged compound with the laser wavelength and/or a more intense laser source, larger responses per pulse are possible. Note that this could also lead to a larger heat artifact. The control points are recorded before the flash when no change is expected.*

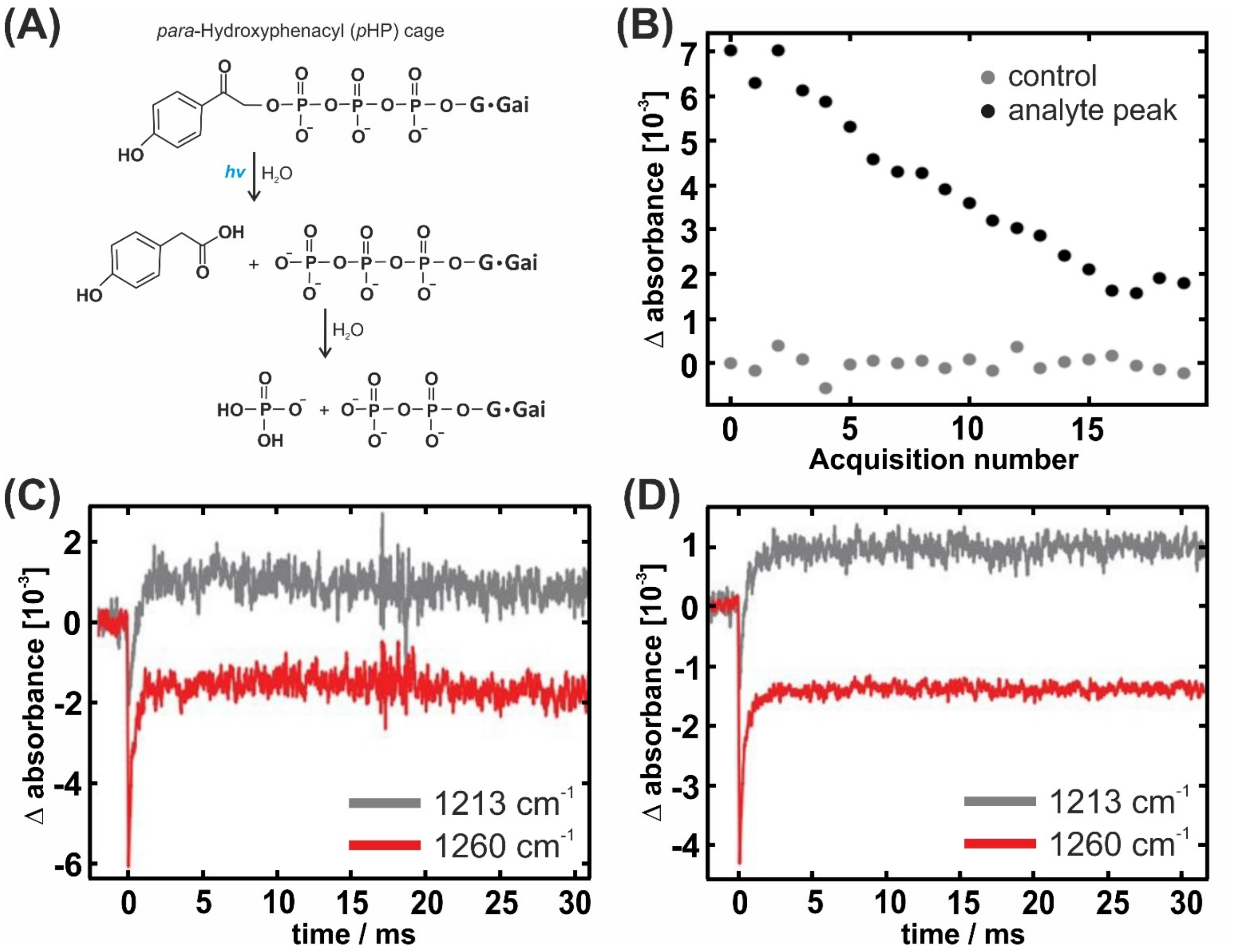


*Figure S4. (A) Photolysis and Hydrolysis of pHP-GTP Reaction Scheme. (B) The magnitude of the spectral response to excitation as a function of excitation pulses. After each flash, the amount of NPE-GTP decreases, leading to a decrease of signal after each pulse. After about 20 pulses all NPE-GTP is consumed. With a better overlap of absorption spectrum of the caged compound with the laser wavelength and/or a more intense laser source, larger responses per pulse are possible. Note that this could also lead to a larger heat artifact. The control points are recorded before the flash when no change is expected. Photolysis spectra obtained by FTIR (black) and dual-comb (red). (C) Kinetics of the hydrolysis reaction obtained by dual-comb experiments. (C) Time resolved absorption change resulted from the first excitation and (D) the average of the first ten experiments at wavenumbers 1213 cm$^{-1}$ and 1260 cm$^{-1}$. Thus (C) shows the S/N per single shot and (D) the S/N per single sample.*

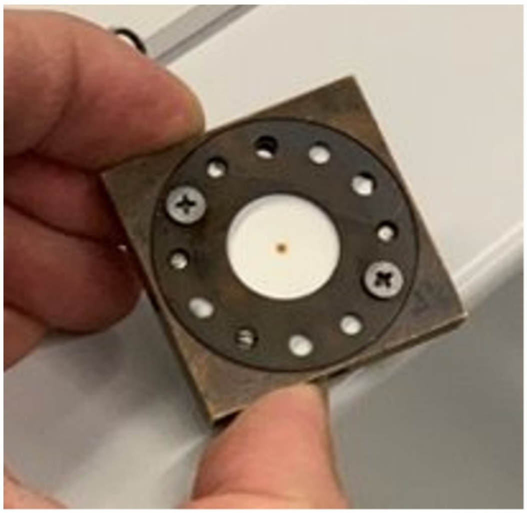

*Figure S5. IR cuvette with photo-sensitive paper, exposed to the QCL laser beam (burned point in the middle). The much smaller diameter of the laser beam in comparison with the about 6mm diameter of the conventionally applied globar light in FTIR offers the advantage of reducing sample consumption by a factor of about 10.*
